# Systematic analysis of REBASE identifies numerous Type I restriction-modification systems that contain duplicated, variable *hsdS* specificity genes that randomly switch methyltransferase specificity by recombination

**DOI:** 10.1101/2020.06.18.137471

**Authors:** John M. Atack, Chengying Guo, Thomas Litfin, Long Yang, Patrick J. Blackall, Yaoqi Zhou, Michael P. Jennings

## Abstract

*N*^6^-adenine DNA methyltransferases associated with some Type I and Type III restriction-modification (R-M) systems are able to randomly switch expression by variation in the length of locus-encoded simple sequence repeats (SSRs). SSR tract-length variation causes ON/OFF switching of methyltransferase expression, resulting in genome-wide methylation differences, and global changes in gene expression. These epigenetic regulatory systems are called phasevarions, phase-variable regulons, and are widespread in bacteria. A distinct switching system has also been described in Type I R-M systems, based on recombination-driven changes in *hsdS* genes, which dictate the DNA target site. In order to determine the prevalence of recombination-driven phasevarions, we generated a program called RecombinationRepeatSearch to interrogate REBASE and identify the presence and number of inverted repeats of *hsdS* downstream of Type I R-M loci. We report that 5.9% of Type I R-M systems have duplicated variable *hsdS* genes containing inverted repeats capable of phase-variation. We report the presence of these systems in the major pathogens *Enterococcus faecalis* and *Listeria monocytogenes*, which will have important implications for pathogenesis and vaccine development. These data suggest that in addition to SSR-driven phasevarions, many bacteria have independently evolved phase-variable Type I R-M systems via recombination between multiple, variable *hsdS* genes.

**Importance:** Many bacterial species contain DNA methyltransferases that have random on/off switching of expression. These systems called phasevarions (phase-variable regulons) control the expression of multiple genes by global methylation changes. In every previously characterised phasevarion, genes involved in pathobiology, antibiotic resistance, and potential vaccine candidates are randomly varied in their expression, commensurate with methyltransferase switching. A systematic study to determine the extent of phasevarions controlled by invertible Type I R-M systems has never before been performed. Understanding how bacteria regulate genes is key to the study of physiology, virulence, and vaccine development; therefore it is critical to identify and characterize phase-variable methyltransferases controlling phasevarions.

## Introduction

Phase variation is the high frequency, random and reversible switching of gene expression (1). Many host-adapted bacterial pathogens posses surface features such as iron acquisition systems (2, 3), pili (4), adhesins (5, 6), and lipooligosaccharide (7, 8) that undergo phase-variable ON-OFF switching of expression by variation in the length of locus encoded simple sequence repeats (SSRs) (1). Variations in SSRs result in the encoded gene being in-frame and expressed (ON), or due to a frameshift downstream of the SSR tract, out-of-frame and not expressed (OFF). Several bacterial pathogens also contain well characterised cytoplasmic *N*^6^-adenine DNA methyltransferases, that are part of restriction-modification (R-M) systems, that exhibit phase-variable expression. We recently characterised the distribution of SSR tracts in Type III *mod* genes and Type I *hsdS, hsdM*, and *hsdR* genes in the REBASE database of restriction-modification (R-M) systems, and demonstrated that 17.4% of all Type III *mod* genes (9), and 10% of all Type I R-M systems contain SSRs that are capable of undergoing phase-variable expression. Phase variation of methyltransferase expression leads to genome-wide methylation differences, which can result in differential regulation of multiple genes in systems known as phasevarions (<u>phase-vari</u>able regul<u>on</u>). Phasevarions controlled by ON-OFF switching of Type III *mod* genes has been well-characterised in a number of host-adapted bacterial pathogens, such as *Haemophilus influenzae* (10, 11), *Neisseria* spp. (12), *Helicobacter pylori* (13), *Moraxella catarrhalis* (14, 15), and *Kingella kingae* (16) (reviewed in (17)). Although we have recently demonstrated that almost 10% of Type I R-M systems contain SSRs, and can potentially undergo phase variation, to-date phase-variable expression of Type I R-M systems has only been demonstrated in two species: an *hsdM* gene switches ON-OFF via SSRs changes in non-typeable *Haemophilus influenzae* (NTHi) (7, 18), and an *hsdS* gene phase varies due to SSRs alterations in *Neisseria gonorrhoeae* (19). The *hsdS* gene in *N gonorrhoeae*, encoding the NgoAV Type I system, contains a G_[n]_ SSR tract, with variation in the length of this tract resulting in either a full length or a truncated HsdS protein being produced, rather than an ON-OFF switch seen with the *hsdM* gene in NTHi and Type III *mod* genes. The full length and truncated HsdS proteins produced from phase variation of the NgoAV system have differing methyltransferase specificities (19).

Type I *hsdS* genes can also undergo phase-variation by recombination between inverted repeats (IRs) encoded in multiple variable copies of *hsdS* genes encoded in the Type I R-M locus (20) and reviewed in (21) (Figure 1A). These systems have been named ‘inverting’ Type I loci, as they phase-vary via ‘inversions’ between the IRs located in the multiple variable *hsdS* genes. The generation of sequence variation by shuffling between multiple protein variants through inverted repeat recombination is perhaps best studied in *pilE* gene encoding pili in *N. gonorrhoeae* (22, 23) and *N. meningitidis* (24). In these systems recombination between a single expressed locus, *pilE*, and multiple adjacent, silent copies of the gene, *pilS*, generate PilE pilin subunit proteins with distinct amino acid sequences. In Type I R-M systems, each HsdS specificity protein is made up of two ‘half’ Target Recognition Domains (TRDs), with each TRD contributing half to the overall specificity of the HsdS protein (Figure 1A). Therefore, changing a single TRD coding region will change the overall specificity of the encoded HsdS protein. The first example of a phasevarion controlled by an inverting Type I R-M system was described in the major human pathogen *Streptococcus pneumoniae* strain D39 (20), and subsequent studies have been conducted in strain TIGR4 (25). This system contains multiple variable *hsdS* loci with inverted repeats, and a locus encoded recombinase, and switches between six alternate HsdS proteins that encode six different methyltransferase specificities (20), and control six different phasevarions. We recently demonstrated the presence of an inverting Type I R-M system in *Streptococcus suis* that switches expression between four alternate HsdS subunits (26). The presence of other inverting Type I systems containing multiple variable *hsdS* genes has also been observed *ad hoc* in several bacterial species, including *Porphyromonas gingivalis* and *Tannerella forsythia* (21, 27). In this study, we carried out a systematic study of the ‘gold-standard’ restriction enzyme database REBASE using a purpose-designed program to systematically identify inverted repeats in *hsdS* genes in order to determine the prevalence of inverting Type I systems in the bacterial domain.

**Figure 1.**
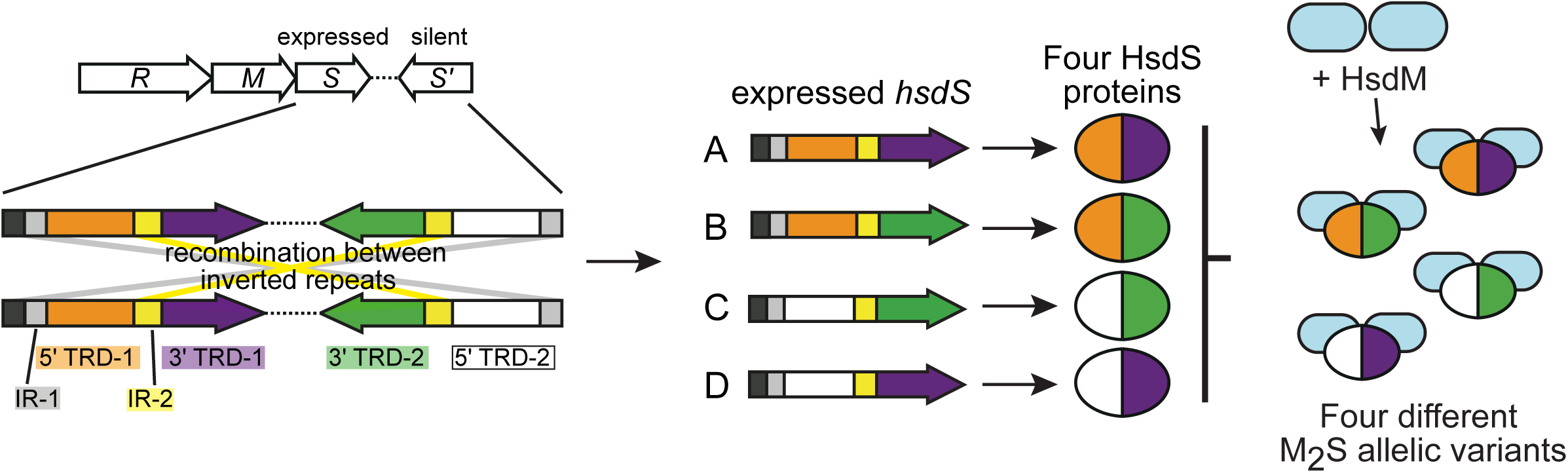
Illustration of how phase-variable switching of inverting Type I systems occurs. Type I R-M loci are made up of three genes, encoding a restriction enzyme (*hsdR; R*), a methyltransferase (*hsdM; M*) and a target sequence specificity protein (*hsdS; S*). Inverting type I systems contain an extra *hsdS* gene termed *hsdS′* (S′). Each *hsdS* gene is made up of two Target Recognition Domains (TRDs). In inverting systems there are multiple variable TRDs present in the two *hsdS* loci. In the illustrated example, there are two different 5′-TRDs (5′-TRD-1 in orange and 5′-TRD-2 in white) and two different 3′ TRDs (3′-TRD-1 in purple and 3′-TRD-2 in green). Inverted repeats are located before 5′-TRD (grey) and between the 5′-TRD and 3′-TRD (yellow). Recombination between these inverted repeats means that four possible *hsdS* coding sequences are present in the expressed *hsdS* locus: allele A = 5′-TRD-1 + 3′-TRD-1; allele B = 5′-TRD-1 + 3′-TRD-2; allele C = 5′-TRD-2 + 3′-TRD-2; allele D = 5′-TRD-2 + 3′-TRD-1. These four different *hsdS* variants mean four different HsdS proteins are produced. Following oligomerisation with an HsdM dimer to form an active methyltransferase, the four different HsdS protein subunits result in four different methyltransferase specificities. This would be described as a ‘four-way’ or ‘four-phase’ switch, as four different HsdS proteins are produced from the four different *hsdS* genes possible in the expressed *hsdS* locus.

## Results

### A systematic search of REBASE reveals that approximately 6% of all Type I R-M systems contain duplicated hsdS loci containing inverted repeats

In order to identify all Type I *hsdS* genes containing inverted repeats (IRs), we searched the restriction enzyme database, REBASE (33), for *hsdS* genes, then searched within 30kb of the start and end of the annotated *hsdS* for inverted repeats (IRs) matching a region of the *hsdS* gene being analysed (see Figure 2). Using the 22,107 *hsdS* genes annotated in REBASE (Supplementary Data 1), we show that 3683 of these *hsdS* genes contain at least one ≥ 20bp sequence with 100% identity to a region that is inverted (i.e., an inverted repeat) and within 30kb of the *hsdS* gene under analysis (Supplementary Data 2). We strictly set our criteria to only select inverted repeats that were 100% identical, and of a minimum size of 20bp in length. This rationale was based on the SpnD39III system, which we described in 2014 (20). The SpnD39III locus contains three different IR regions that are 15bp, 85bp, and 33bp long, encoded within multiple variable *hsdS* genes. Therefore, setting our minimum length criteria for an IR at 20bp means any IRs detected are above the length shown previously to result in homologous recombination between variable *hsdS* genes.

**Figure 2.**
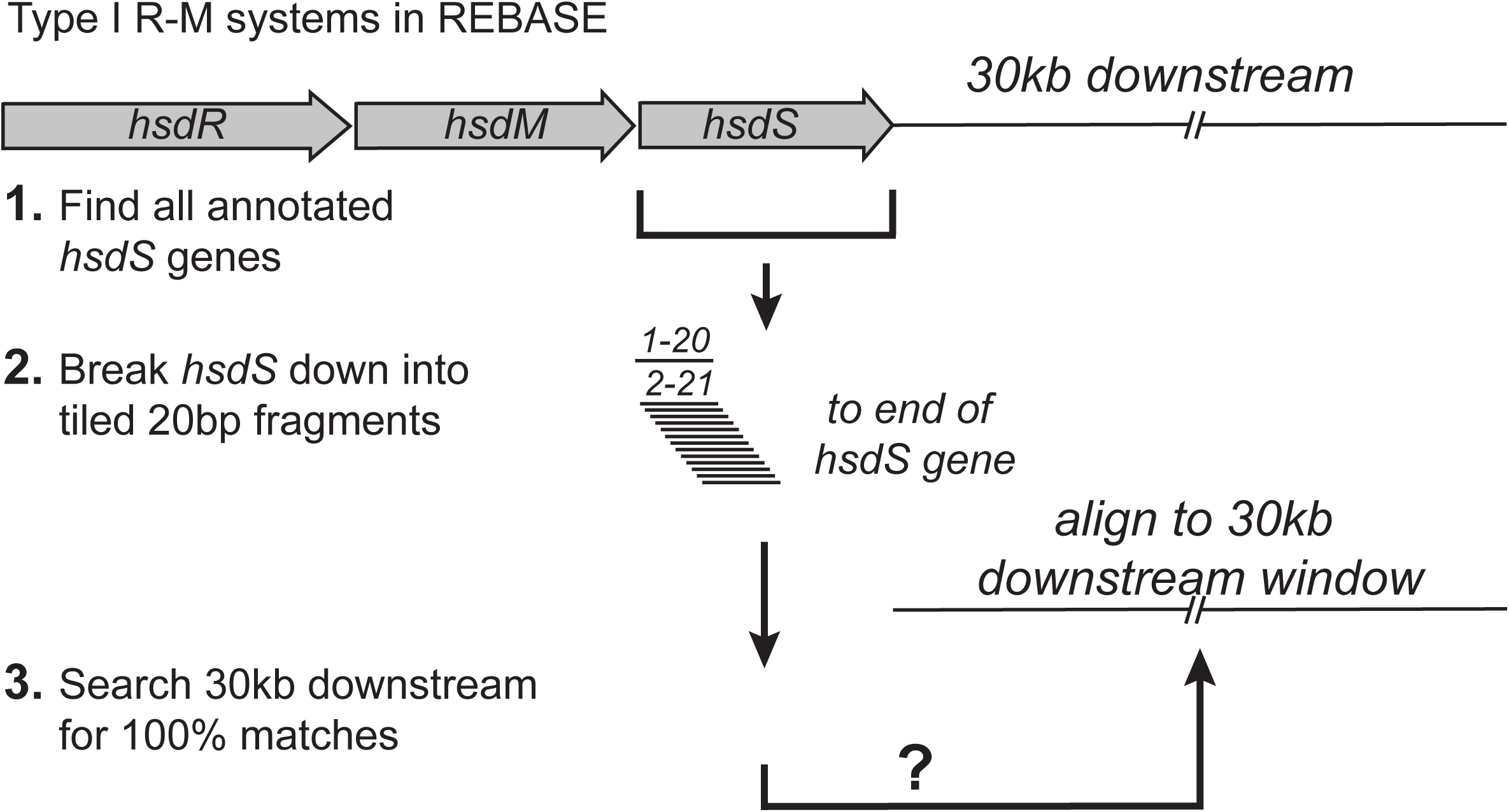
Illustration of our search methodology. All Type I *hsdS* loci were downloaded from REBASE. These loci were then broken down into 20bp tiled fragments, each staggered by 1 bp (fragment 1 = bp1-20, fragment 2 = bp2-21, etc). These tiles were then used as a search term to search for 100% identical fragments in the opposite orientation, i.e., inverted, 30kb upstream of the annotated start codon and 30kb downstream of the annotated stop codon of the *hsdS* gene under investigation. Although we searched both upstream and downstream of the annotated *hsdS* gene understudy, we have only shown the downstream search in this illustration for simplicity.

We carried out our search for inverted repeats using a bespoke perl script (irepeat.upstream.pl), which we have made available at https://github.com/GuoChengying-7824/type_I. This script was also implemented as a simple, easy-to-use server called ‘RecombinationRepeatSearch’, which can be found at https://sparks-lab.org/server/recombinationrepeatsearch/. This software allows a user to input any gene or DNA sequence (e.g., an *hsdS* gene) and by providing the relevant upstream and downstream DNA sequence (e.g., the *hsdS* gene plus 30kb upstream and downstream as a single sequence), the software is able to locate regions containing inverted repeats (see Figure 2).

Our analysis showed that of the 3683 *hsdS* genes containing at least one IR, many *hsdS* genes had more than one downstream IR, and so were counted twice (for an *hsdS* gene with two downstream inverted repeats), three times (for an *hsdS* gene with three downstream inverted repeats), and so on. Therefore, in order to determine the number of individual *hsdS* genes with at least one downstream IR, we collated together all identical *hsdS* genes. Followin this collation, we show that 991 individual Type I R-M loci have *hsdS* genes with *at least* one IR located within 30kb (Supplementary Data 3). Taking into account all bacterial strains with at least one full Type I R-M system (at least one *hsdR, hsdM* and *hsdS* gene; 14830 strains in total) and where the IR(s) are in a second, duplicated *hsdS* within the same Type I R-M locus, 875 contain at least one IR in a second, duplicated, variable *hsdS* gene within the same Type I locus. This equates to 5.9% (875/14830) of all Type I R-M systems being potentially phase-variable, and therefore able to control phasevarions.

Our analysis shows that some bacterial species contain a relatively low proportion of examples of strains that have IRs within 30kb of annotated *hsdS* genes. For example, there are 428 *Staphylococcus aureus* genomes in REBASE, and of these, only 5 contain an *hsdS* gene with an IR located within 30kb (Supplementary Data 3); of the 232 *Pseudomonas aeruginosa* genomes examined, only 1 contained an *hsdS* with an IR found within 30kb. Detailed analysis of these regions revealed that the IR found within 30kb of the annotated *hsdS* gene in *P. aeruginosa* strain SPA01 (accession number LQBU01000001) is only 28bp long, and although it is possible that inversions do occur between these inverted repeats, the IR is not in a locus annotated as an *hsdS*. Manual examination of the 5 IRs found within 30kb of annotated *hsdS* genes in *S. aureus* also do not appear in a second annotated *hsdS* locus. Three of these inverted repeats in *S. aureus* are >200bp long (in strains 333, M013, and UCI 48); for example, the IR found within 30kb of the *hsdS* annotated as S.SauM013ORF1818P in *S. aureus* strain M013 (accession number CP003166; Supplementary Data 1 & 2) is 529bp long. The S.SauM013ORF1818P locus is itself 531bp long. It is likely that these two regions are able to recombine, and flank a region including genes for a hyaluronate lyase and a metalloproteinase. It was recently demonstrated in *S. aureus* that recombination between two Type I loci approximately 1.26Mb apart are able to mediate genome inversions (34). It is therefore possible that a small proportion of the large (>200bp) IRs we identified in our search (Supplementary Data 2) are part of larger inverting DNA segments, and not associated with individual Type I loci that undergo rearrangements between expressed and silent *hsdS* genes contained in a single Type I locus, i.e, not part of inverting Type I R-M systems.

Using the SpnD39III system present in *S. pneumoniae*, which we identified as the first inverting, phase-variable Type I R-M system, and the first example of a phasevarion in a Gram-positive bacterium (20), we show that of the 78 *S. pneumoniae* strains listed in REBASE, all of the strains where we were able to obtain the annotated genome (52 total) contain the SpnD39III system. This confirmed the findings in our 2014 study, where we showed every genome in GenBank (n=262) contained a Type I locus where inverted variable *hsdS* genes were present (20). Our systematic search of REBASE also identified the Type I system in *S. suis* which we have previously shown to shuffle between four different HsdS proteins (26). These findings serve as a ‘positive control’ for our search methodology, in that it is able to identify systems previously shown to contain IRs and to be phase-variable by *ad-hoc* searches.

Our search confirms the presence of inverting Type I R-M systems with downstream IRs identified previously. For example, we show that 7 out of 15 strains of *P. gingivalis* with an annotated genome in REBASE contain *hsdS* genes with IRs located within 30kb, and 2 out of 7 strains of *T. forsythia* contain annotated *hsdS* genes where IRs are present within 30kb (27). Our analysis of these regions confirmed the IRs to be present in a second, variable *hsdS* gene that is part of the same Type I R-M locus, and which we class as an inverting, i.e., a phase-variable Type I locus. Using these systems as an example, and based on previous work with the SpnIII system in *S. pneumoniae* (20), and the inverting Type I system in *S. suis* (26), we analysed the regions immediately upstream of both *hsdS* genes present in each individual *P. gingivalis* and *T. forsythia* Type I locus containing IRs. This analysis demonstrated that only the *hsdS* gene immediately downstream of the *hsdM* gene is a functional open-reading frame, with the second downstream *hsdS* gene encoded on the opposite strand being silent (*hsdS*′), as this second gene does not contain an ATG start codon or a region recognised as a promoter using the bacterial promoter prediction tools CNNpromoter_b (35) and PePPER (36).

### Three major veterinary pathogens contain Type I R-M systems containing duplicated variable hsdS loci

Many species contained a high prevalence strains with *hsdS* genes with downstream IRs, and with these IRs located within a separate, variable *hsdS* genes that were part of the same Type I locus containing the *hsdS* gene under study. For example, we identified Type I R-M systems with multiple *hsdS* genes in two major veterinary pathogens, in addition to the one identified in *S. suis* (Figure 3A; Supplementary Data 3). In the pig pathogen, *Actinobacillus pleuropneumoniae*, of the 23 genomes available in REBASE, 18 contain at least one Type I R-M system with multiple, variable inverted *hsdS* loci, and with these *hsdS* genes containing the IRs identified by our search. In the cattle pathogen *Mannheimia haemolytica*, 19 out of the 23 strains surveyed contain at least one Type I R-M system with multiple, variable inverted *hsdS* loci with IRs. Detailed examination of each of the inverting Type I R-M systems we identified in *A. pleuropneumoniae* and *M. haemolytica* showed that these systems also contain a gene encoding a recombinase/integrase, and additional genes encoding proteins unknown function (Figure 3A). In addition, our survey demonstrated that 24 out of 42 *S. suis* strains analysed contain an inverting Type I system, confirming our earlier observation that the Type I system in this species is not present in all strains, but conserved within a virulent lineage that causes zoonotic infections (26). In all three of these veterinary pathogens, two IRs are present in a second distinct *hsdS* gene (*hsdS*′) immediately downstream of the *hsdS* understudy, and part of the same Type I R-M locus (Figure 1). Examination of the location of each pair of IRs present in these two *hsdS* genes demonstrated they occurupstream of the 5′-TRD, and between the 5′-TRD and 3′-TRD (Figure 1, Figure 3). The presence of multiple IRs that are in a second variable *hsdS* gene (*hsdS*′) immediately downstream of the *hsdS* gene under study is highly indicative that these *hsdS* genes undergo inversions, i.e., they are phase-variable.

**Figure 3.**
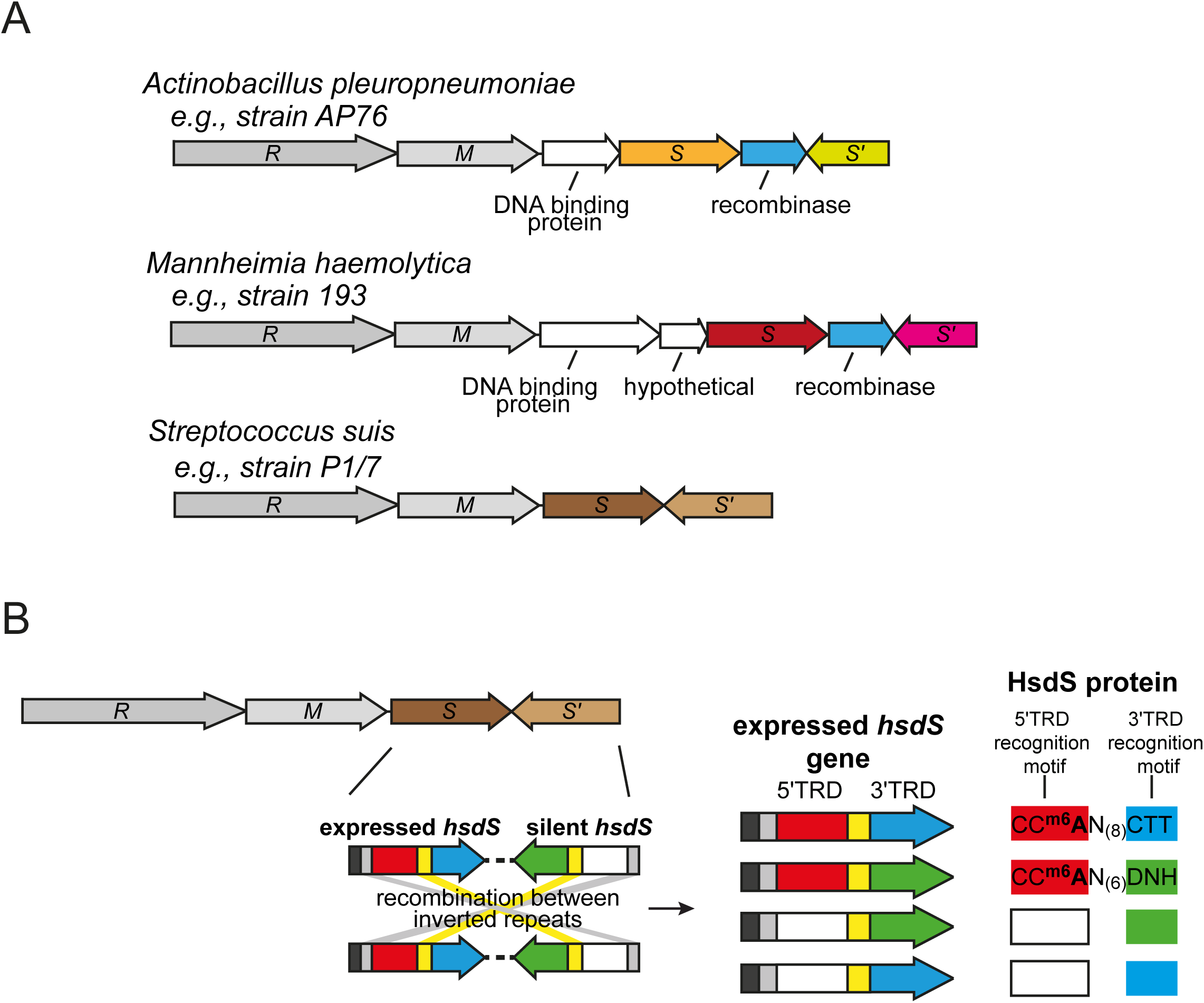
**A) schematic representation of Type I loci with multiple variable *hsdS* genes containing inverted repeats from three important veterinary pathogens.** Coloured arrows represent variable *hsdS* genes. Blue arrows indicate that a gene with high identity to a recombinase/integrase is present at the locus**; B) Illustration of the mode of switching of the four-way switch occurring in *Streptococcus suis***. *S. suis* contains a Type I locus containing duplicated variable *hsdS* loci containing inverted repeats (SSU1271-SSU1274 in *S. suis* strain P1/7). As illustrated in Figure 1, each *hsdS* gene is made up of separate 5′ (red and white) and 3′ (blue and green) TRDs. Inverted repeats are present before the 5′ TRD (grey) and between the 5′ and 3′ TRDs (yellow). Each TRD recognises a different 3bp DNA sequence, giving rise to 4 separate HsdS proteins that are predicted to methylate four different DNA sequences dependent on the TRDs present. We have solved the specifity of allele A (5′TRD-1 [red] + 3′TRD-1 [blue]) and allele B (5′TRD-1 [red] + 3′TRD-2 [green]). 5′TRD-1 (red) recognises CCA, 3′TRD-1 (blue) recognises CTT, 3′TRD-2 (green) recognises DNH. D = A, G, or T; N = any nucleotide; H = A, C, or T. XXX = the recognition motif is undetermined.

We cloned and over-expressed two *hsdS* alleles, alleles A and B, of the Type I inverting system that we found in *S. suis* (26) in order to solve the methyltransferase specificity of the Type I methyltransferases containing these HsdS proteins. We have used this approach extensively with Type III *mod* genes in order to solve specificity (5, 9), with the same site observed using the native protein using genomic DNA from the actual species and the over-expressed protein in *E. coli* (26). We only expressed HsdS alleles A and B as we do not observe any strains of *S. suis* with annotated genomes where either allele C or allele D (Figure 3B) is present in the *hsdS* expressed locus immediately downstream of the *hsdM* (26). This approach demonstrated that allele A methylates the sequence CC^m6^AN_(8)_CTT, and allele B methylates the sequence CC^m6^AN_(6)_DNH (D = A, G, or T; H = A, C, or T; N = any nucleotide). This is consistent with allele A and allele B sharing the same 5′-TRD (giving the same half recognition sequence of CCA), but a different 3′-TRD (giving different half recognition sequences of CTT, and DNH, respectively) (Figure 3B). Solving the specificity of the two most common alleles found in the expressed *hsdS* locus of this phase-variable system (26) provides valuable information required to fully characterise the gene expression differences that result from the phase-variation of this system.

### The major human and veterinary pathogen Listeria monocytogenes contains an inverting Type I R-M system that appears to be associated with virulence

Our analysis shows that an inverting Type I R-M system is present in approximately half of all strains of *Listeria monocytogenes* that are deposited in REBASE (60 out of 123 strains). This inverting Type I system was previously identified in *L. monocytogenes* ST8 strains associated with disease in aquaculture and poultry farming (21, 37). Different *hsdS* sequences are present in the expressed *hsdS* locus of multiple strains of *L. monocytogene*s (37), although no recombination has been demonstrated within an individual strain. Phylogenetic analysis of these strains (Figure 4) shows that strains containing this system cluster in specific clades. This data suggests that selection and expansion of strains containing this system is occurring, with a possible association between this system and with strains that persist in fish and chickens (37). Analysis of the phenotypes regulated by this system may have an impact on vaccine and pathogenesis studies of this important human and veterinary pathogen.

**Figure 4.**
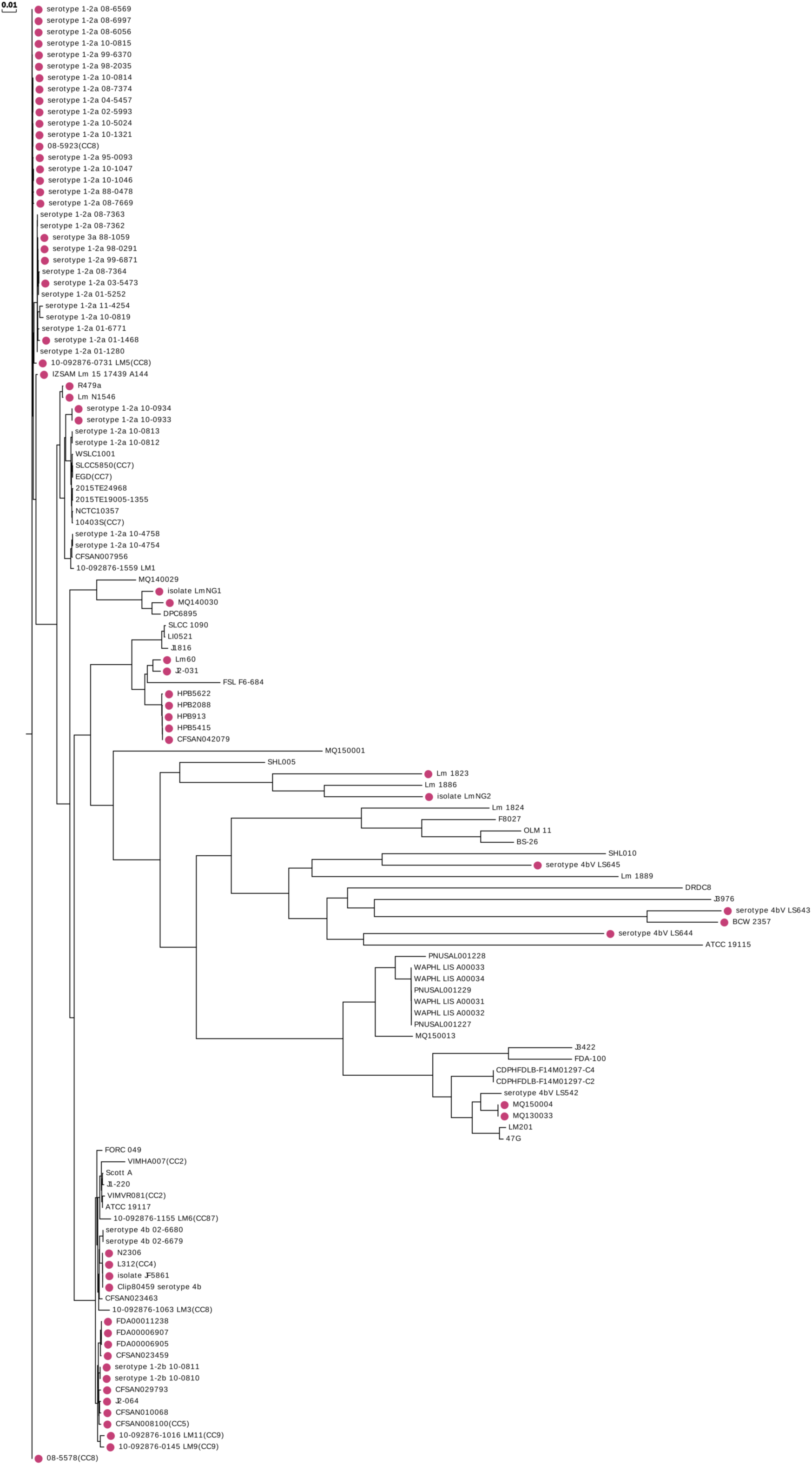
The whole-genome phylogenic tree was constructed by CVTree (Version-3.0.0) for 128 strains of *Listeria monocytogenes* annotated in REBASE. Red circles indicate strains that containing Type I systems containing duplicated *hsdS* genes containing inverted repeats. The distance measures the dissimilarity of each strain.

### The nosocomial, antibiotic-resistant pathogen Enterococcus faecalis contains a highly diverse phase variable Type I R-M locus that is widely distributed

We identify a Type I R-M system containing multiple variable *hsdS* loci containing IRs present in *Enterococcus faecalis*, a multidrug-resistant, nosocomial pathogen of major medical importance. This system has been previously noted to occur in a single strain of *E. faecalis* (21), but no systematic study of the distribution of this system in *E. faecalis* had been carried out. This system is present in 24 out of the 34 strains of *E. faecalis* present in REBASE. Analysis of the sequences of each of the 24 Type I loci containing duplicated *hsdS* genes (Figure 5A) shows a high level of variability at each individual *hsdS* locus, with thirteen different 5′-TRDs, and sixteen different 3′-TRDs present in the *hsdS* genes annotated in REBASE. This data is highly indicative of shuffling of TRDs, and shows significant inter-strain variability. Our phylogenetic analysis of the strains of *E. faecalis* containing this system (Figure 5B) shows that the presence of the Type I R-M system is widely distributed within the overall *E. faecalis* population, and not associated with a particular lineage or groups of strains. This inverting Type I R-M locus also contains an integrase/recombinase, in addition to multiple variable *hsdS* genes containing IRs, adding further weight to the evidence that this system is phase-variable.

**Figure 5.**
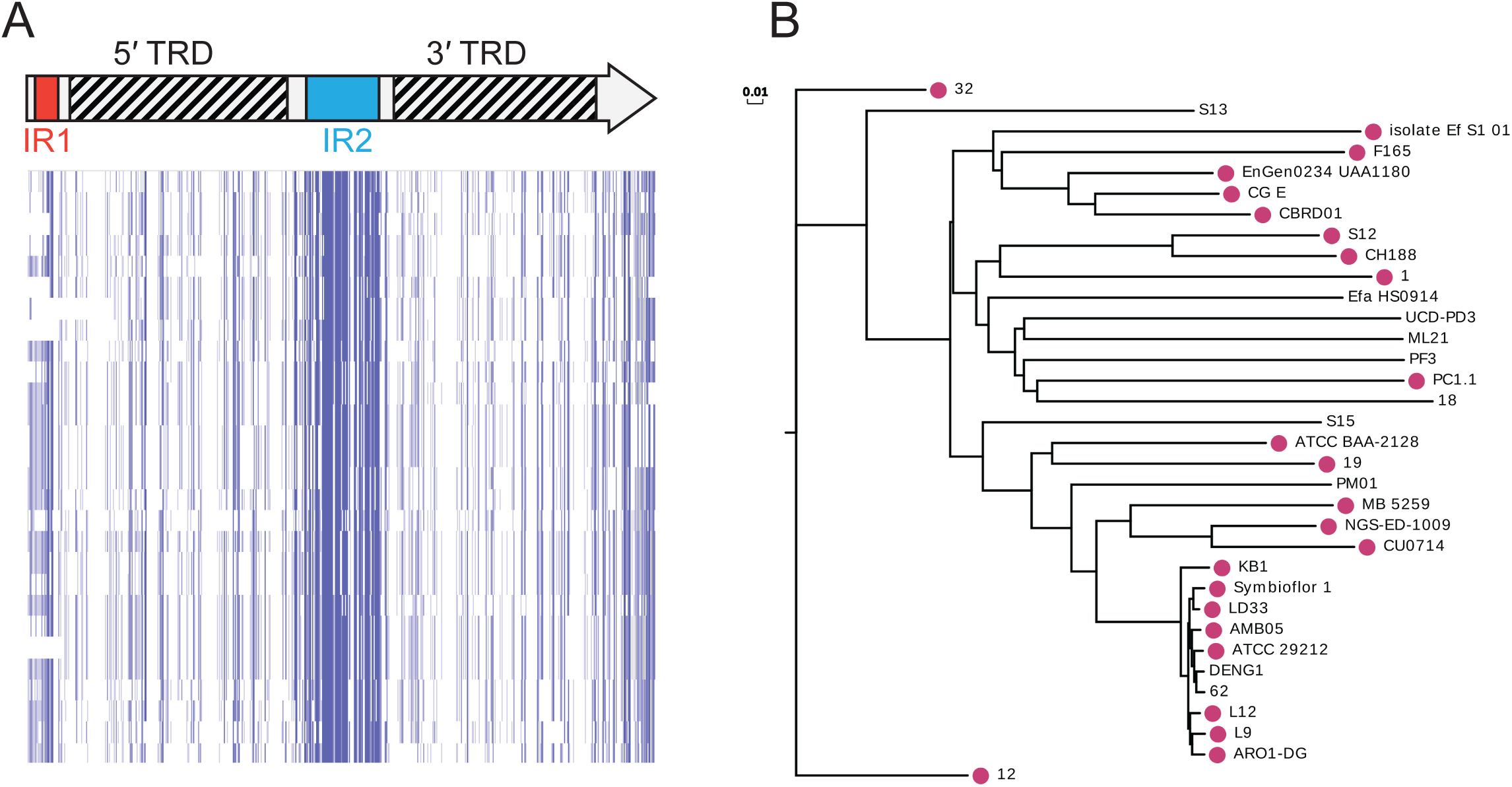
**A)** Type I *hsdS* gene showing the location of the 5′ and the 3′ TRD, and the inverted repeats. Sequence analysis of representative examples of each *hsdS* gene present in *Enterococcus faecalis*. Alignments were carried out using ClustalW, and visualized in JalView overview feature. Blue colour indicates % nucleotide identity; **B)** The whole genome phylogenic tree was constructed by CVTree (Version-3.0.0) for 34 strains of *Enterococcus faecalis* annotated in REBASE. Red circles indicate strains that containing Type I systems containing duplicated *hsdS* genes containing inverted repeats. The distance measures the dissimilarity of each strain.

## Discussion

This is the first time, to our knowledge, that a systematic study has been carried out to identify Type I R-M systems that contain inverted repeats that are capable of mediating phase-variable expression, and thereby potentially control phasevarions. A previous study demonstrated that integrases/recombinases with high homology to the integrase present in the SpnD39III locus (20) were widespread in the bacterial domain (21). In order to carry out our systematic analysis, we designed software to specifically search for inverted repeats in DNA (code available at https://github.com/GuoChengying-7824/type_I), and applied strict selection criteria so that we only identified inverted DNA repeats that are longer than those that have previously been shown to result in homologous recombination between variable *hsdS* genes (20). We limited the distance away from the *hsdS* locus understudy (30kb) in order to only identify distinct ‘inverting’ Type I R-M systems. We have made this software available as a user-friendly server (RecombinationRepeatSearch; https://sparks-lab.org/server/recombinationrepeatsearch/), which allows the user to search any DNA sequence for inverted repeat regions.

By limiting our selection criteria (100% IR identity; minimum IR length of 20bp; 30kb window upstream and downstream each *hsdS*), we have likely missed some Type I loci that are ‘inverting’; for example, we will miss any IRs that are <20bp, and we would not detect any *hsdS* containing IRs that are over 30kb away. However, we would argue that *hsdS* genes located over 30kb away from each other would not comprise a single ‘inverting’ Type I *hsd* locus, and that the recombination of these separate *hsdS* genes may not control phasevarions. We also identified a small number of large (>200bp) IRs present within 30kb of annotated *hsdS* genes, but a manual examination of these systems revealed that the IRs are not present in a second *hsdS* gene.

Our systematic analysis of REBASE identified Type I loci containing multiple *hsdS* genes where we detect IRs in a range of commensal organisms such as *Bacteroides fragilis* and multiple *Ruminococcus* species, in environmental bacterial species such as *Leuconostoc mesenteroides*, and in a number of *Lactobacillus* species that are important to the biotechnology and food production imdustries (Supplementary Data 3). This reflects our previous studies where we observed simple sequence repeats that mediate phase-variation in multiple Type I (38) and Type III methyltransferase genes (9) present in a variety of commensal and environmental organisms. One obvious reason for generating diversity in methyltransferase specificity is that it will increase resistance to bacteriophage. However, in every case where a methyltransferase has been demonstrated to phase-vary, it has also been shown to comprise a phasevarion; therefore in addition to improving survival when exposed to bacteriophage, phase-variable methyltransferases are also likely to increase the phenotypic diversity present in a bacterial population, providing bacteria that encode them an extra contingency strategy to deal with changing environmental conditions. It will be interesting to determine how such plasticity of gene expression would be advantageous in a changing environment that cannot be dealt with via conventional “sense and respond” gene regulation strategies (1), particularly as regards phage resistance.

We identified multiple variable *hsdS* loci that contain IRs in the major human pathogens *L. monocytogenes* and *E. faecalis*. Our analysis also demonstrated that a variety of veterinary pathogens, contain Type I systems where IRs are present in multiple variable *hsdS* genes. Many of the veterinary pathogens that we show contain inverting Type I loci also contain separate, distinct Type III or Type I R-M systems that are capable of phase-varying via changes in locus located simple sequence repeats. These species include *Actinobacillus pleuropneumoniae, Mannheimia haemolytica, Streptococcus suis, Haemophilus (Glasserella) parasuis*, and multiple *Mycoplasma* species (9, 38). This means all these veterinary pathogens have evolved phase-variation of both Type I and Type III methyltransferases, and in the case of Type I systems, by both SSR tract length changes (38) and by recombination between variable *hsdS* genes containing IRs (this study). For example, *A. pleuropneumoniae* encodes two distinct Type III methyltransferase (*mod*) genes containing simple sequence repeats (9), and a Type I system containing variable *hsdS* loci where IRs are present (this study; Figure 3A). We predict that this inverting Type I system switches between four separate *hsdS* genes, and therefore results in four different methyltransferase specificities. This means that there are a total of sixteen different combinations of methyltransferase activity potentially present in a population of *A. pleuropneumoniae*. Therefore it is critical to determine the genes and proteins that are part of the phasevarions in these species, although this will not be a simple task due the breadth and diversity of the variable methyltransferases present in these organisms.

In summary, we identify that 5.9% of Type I R-M systems contain duplicated variable *hsdS* genes containing inverted repeats, are likely to phase vary, and consequently comprise a phasevarion. A broad range of bacterial species encode these systems. Our previous work showed that 2% of Type I *hsdM* and 7.9% of Type I *hsdS* genes contain SSRs (38). Together with our findings in this study, this means that 15.8% of all Type I systems are capable of phase-variable expression. In addition, previous studies have shown that 17.4% of Type III methyltransferases contain SSRs (9) and therefore capable of phase-varying. That approximately the same percentage of two independent DNA methyltransferase systems have evolved the ability to phase-vary in expression demonstrates that generating variation via switching of methyltransferase expression is a widespread strategy used by bacteria, and that this method of increasing diversity has evolved independently multiple times. The study of phasevarions is not only key to vaccine development against pathogenic bacteria that contain them, but necessary to understand gene expression and regulation in the bacterial domain.

## Materials and Methods

### REBASE survey and bioinformatics

All gene sequences of Type I *hsdS* subunits were downloaded from http://rebase.neb.com/rebase/rebase.seqs.html. The annotation for each gene was downloaded from http://rebase.neb.com/rebase/rebadvsearch.html. A total of 22,107 genes were obtained with complete annotation information, which includes the start, end, and genomic information of the gene. However, the annotation does not contain the information regarding if the gene is in the positive or the negative strand of the genome. This information is obtained after aligning the gene sequence with the corresponding genomic sequence. All genomic sequences were downloaded from NCBI GenBank, and a total of 15,486 genomes were downloaded. After a gene is located in the corresponding genome, we obtained both 30kb upstream of the annotated start codon and 30kb downstream of the annotated stop codon. The 30kb upstream and downstream regions were compared against 20-500 bp fragments of the reverse gene sequence. No reverse search is performed if a gene is in the negative strand. If upstream and downstream regions contain a region mapping to a 500 bp reverse fragment, we further scanned the fragment length between 500 and 1500 bp. This process is implemented by a perl script (irepeat.upstream.pl) located at https://github.com/GuoChengying-7824/type_I. We also established this software as a server called RecombinationRepeatSearch, and is located at https://sparks-lab.org/server/recombinationrepeatsearch/. This allows a user to input their gene of interest, and by including the respective upstream or downstream genomic sequence, they are able to determine if the DNA sequence of their gene of interest encodes inverted DNA repeats in the immediate vicinity. Following this search, all redundant repeating segments were removed by filtering. Only 100% matches for inverted repeats are recorded. All inverted repeat regions found are listed in Supplementary Data 2. Phylogenetic trees were constructed using the neighboring method (Neighbor-joining) using CVTree (Version-3.0.0) version (28, 29), with the default Hao method, and a K value of 6, as recommended for prokaryotic trees (30).

### Cloning and over-expression of the phase-variable Type I system from Streptococcus suis

The entire *hsdMS* region from *S. suis* strain P1/7 containing *hsdS* allele B was cloned using primers SsuT1-oE-F (5′-AGTCAG CCATGG GG TCA ATT ACA TCA TTT GTT AAA CGA ATA CAA G) and SsuT1-oE-R (5′-AGTCAG GGATCC TCA GTA ATA AAG TTG GGC AAC TTT TTC) into the NcoI-BamHI site of vector pET15b (Novagen). In order to generate *hsdS* allele A, 3′ -TRD allele 1 was synthesised as a gBLOCK (IDT) and cloned into pET15b::allele B that was linearised either side of 3′-TRD allele 2 using primers TRD-Swap-inv-F (5′-CTG CTG CCA CCG CTG AGC AAT AAC TAG C) and TRD-Swap-inv-R (5′-CTT CCC ATA AGG AGA GTT ATC ATC TCC), to generate vector pET15b::allele A. Inverse PCR using this construct was carried out with KOD polymerase (EMD Millipore) according to manufacturers instructions. Following sequencing to confirm constructs were correct, over-expression of each methyltransferase (HsdM plus either HsdS allele A or HsdS allele B) was carried out using *E. coli* BL21 cells, which were induced by the addition of IPTG to a final concentration of 0.5mM over-night at 37°C with shaking at 200rpm. Over-expression was confirmed by SDS-PAGE by comparing to an uninduced control.

### Single-Molecule, Real-Time (SMRT) sequencing and methylome analysis

Genomic DNA from *E. coli* cells expressing the *S. suis* HsdM plus either allele A or allele B HsdS were prepared using the Sigma GenElute genomic DNA kit according to the manufacturer’s instructions. SMRT sequencing and methylome analysis was carried out as previously (31, 32). Briefly, DNA was sheared to an average length of approximately 10-20 kb using g-TUBEs (Covaris; Woburn, MA, USA) and SMRTbell template sequencing libraries were prepared using sheared DNA. DNA was end-repaired, then ligated to hairpin adapters. Incompletely formed SMRTbell templates were degraded with a combination of Exonuclease III (New England Biolabs; Ipswich, MA, USA) and Exonuclease VII (USB; Cleveland, OH, USA). Primer was annealed and samples were sequenced on the PacBio RS II (Menlo Park, CA, USA) using standard protocols for long insert libraries. SMRT sequencing and methylome analysis was carried out by SNPSaurus (University of Oregon, USA).

## Acknowledgements

We thank Eric and Allison from SNPsaurus, University of Oregon, USA, for expert technical assistance in carrying out SMRT sequencing and methylome analysis. This work was supported by the Australian National Health and Medical Research Council (NHMRC) Program Grant 1071659 and Principal Research Fellowship 1138466 to MPJ; Project Grant 1099279 to JMA and 1121629 to YZ; Australian Research Council (ARC) Discovery Projects 170104691 to MPJ, 180100976 to JMA and PJB, and 180102060 to YZ. Funding for open access charge: National Health and Medical Research Council, Australia.

## Figure legends

**Supplementary Data 1** – all Type I *hsdS* genes downloaded from REBASE

**Supplementary Data 2** – all IRs found in *hsd*S genes

**Supplementary Data 3** – all representative *hsdS* genes with IRs

## References

1. Moxon R, Bayliss C, Hood D. 2006. Bacterial contingency loci: the role of simple sequence DNA repeats in bacterial adaptation. Ann Rev Genet 40:307–333.

2. Ren Z, Jin H, Whitby PW, Morton DJ, Stull TL. 1999. Role of CCAA nucleotide repeats in regulation of hemoglobin and hemoglobin-haptoglobin binding protein genes of *Haemophilus influenzae*. J Bacteriol 181:5865–70.

3. Richardson AR, Stojiljkovic I. 1999. HmbR, a hemoglobin-binding outer membrane protein of *Neisseria meningitidis*, undergoes phase variation. J Bacteriol 181:2067–74.

4. Blyn LB, Braaten BA, Low DA. 1990. Regulation of *pap* pilin phase variation by a mechanism involving differential dam methylation states. EMBO J 9:4045–54.

5. Atack JM, Winter LE, Jurcisek JA, Bakaletz LO, Barenkamp SJ, Jennings MP. 2015. Selection and counter-selection of Hia expression reveals a key role for phase-variable expression of this adhesin in infection caused by non-typeable *Haemophilus influenzae*. J Infect Dis 212:645–53.

6. Dawid S, Barenkamp SJ, St. Geme JW. 1999. Variation in expression of the *Haemophilus influenzae* HMW adhesins: A prokaryotic system reminiscent of eukaryotes. Proc Natl Acad Sci U S A 96:1077–1082.

7. Fox KL, Atack JM, Srikhanta YN, Eckert A, Novotny LA, Bakaletz LO, Jennings MP. 2014. Selection for phase variation of LOS biosynthetic genes frequently occurs in progression of non-typeable *Haemophilus influenzae* infection from the nasopharynx to the middle ear of human patients. PLoS One 9:e90505.

8. Poole J, Foster E, Chaloner K, Hunt J, Jennings MP, Bair T, Knudtson K, Christensen E, Munson RS, Jr., Winokur PL, Apicella MA. 2013. Analysis of nontypeable *Haemophilus influenzae* phase variable genes during experimental human nasopharyngeal colonization. J Infect Dis 208:720–727.

9. Atack JM, Yang Y, Seib KL, Zhou Y, Jennings MP. 2018. A survey of Type III restriction-modification systems reveals numerous, novel epigenetic regulators controlling phase-variable regulons; phasevarions. Nucleic Acids Res 46:10.1093/nar/gky192.

10. Atack JM, Srikhanta YN, Fox KL, Jurcisek JA, Brockman KL, Clark TA, Boitano M, Power PM, Jen FEC, McEwan AG, Grimmond SM, Smith AL, Barenkamp SJ, Korlach J, Bakaletz LO, Jennings MP. 2015. A biphasic epigenetic switch controls immunoevasion, virulence and niche adaptation in non-typeable *Haemophilus influenzae*. Nat Commun 6:doi: 10.1038/ncomms8828.

11. Srikhanta YN, Maguire TL, Stacey KJ, Grimmond SM, Jennings MP. 2005. The phasevarion: A genetic system controlling coordinated, random switching of expression of multiple genes. Proc Natl Acad Sci U S A 102:5547–5551.

12. Srikhanta YN, Dowideit SJ, Edwards JL, Falsetta ML, Wu H-J, Harrison OB, Fox KL, Seib KL, Maguire TL, Wang AHJ, Maiden MC, Grimmond SM, Apicella MA, Jennings MP. 2009. Phasevarions mediate random switching of gene expression in pathogenic *Neisseria*. PLoS Pathog 5:e1000400.

13. Srikhanta YN, Gorrell RJ, Steen JA, Gawthorne JA, Kwok T, Grimmond SM, Robins-Browne RM, Jennings MP. 2011. Phasevarion mediated epigenetic gene regulation in *Helicobacter pylori*. PLoS One 6:e27569.

14. Blakeway LV, Power PM, Jen FE, Worboys SR, Boitano M, Clark TA, Korlach J, Bakaletz LO, Jennings MP, Peak IR, Seib KL. 2014. ModM DNA methyltransferase methylome analysis reveals a potential role for *Moraxella catarrhalis* phasevarions in otitis media. FASEB J 28:5197–5207.

15. Seib KL, Peak IR, Jennings MP. 2002. Phase variable restriction-modification systems in *Moraxella catarrhalis*. FEMS Immunol Med Mic 32:159–165.

16. Srikhanta YN, Fung KY, Pollock GL, Bennett-Wood V, Howden BP, Hartland EL. 2017. Phasevarion regulated virulence in the emerging paediatric pathogen *Kingella kingae*. Infect Immun 85:e00319–17.

17. Atack JM, Tan A, Bakaletz LO, Jennings MP, Seib KL. 2018. Phasevarions of Bacterial Pathogens: Methylomics Sheds New Light on Old Enemies. Trends in Microbiology 26:715–726.

18. Zaleski P, Wojciechowski M, Piekarowicz A. 2005. The role of Dam methylation in phase variation of *Haemophilus influenzae* genes involved in defence against phage infection. Microbiology 151:3361–9.

19. Adamczyk-Poplawska M, Lower M, Piekarowicz A. 2011. Deletion of One Nucleotide within the Homonucleotide Tract Present in the hsdS Gene Alters the DNA Sequence Specificity of Type I Restriction-Modification System NgoAV. J Bacteriol 193:6750–6759.

20. Manso AS, Chai MH, Atack JM, Furi L, De Ste Croix M, Haigh R, Trappetti C, Ogunniyi AD, Shewell LK, Boitano M, Clark TA, Korlach J, Blades M, Mirkes E, Gorban AN, Paton JC, Jennings MP, Oggioni MR. 2014. A random six-phase switch regulates pneumococcal virulence via global epigenetic changes. Nat Commun 5:doi: 10.1038/ncomms6055.

21. De Ste Croix M, Vacca I, Kwun MJ, Ralph JD, Bentley SD, Haigh R, Croucher NJ, Oggioni MR. 2017. Phase-variable methylation and epigenetic regulation by type I restriction– modification systems. FEMS Microbiol Rev 41:S3–S15.

22. Helm RA, Seifert HS. 2010. Frequency and rate of pilin antigenic variation of Neisseria meningitidis. J Bacteriol 192:3822–3.

23. Seifert HS. 1996. Questions about gonococcal pilus phase- and antigenic variation. Mol Microbiol 21:433–40.

24. Sechman EV, Rohrer MS, Seifert HS. 2005. A genetic screen identifies genes and sites involved in pilin antigenic variation in Neisseria gonorrhoeae. Mol Microbiol 57:468–83.

25. Oliver MB, Basu Roy A, Kumar R, Lefkowitz EJ, Swords WE. 2017. *Streptococcus pneumoniae* TIGR4 Phase-Locked Opacity Variants Differ in Virulence Phenotypes. mSphere 2.

26. Atack JM, Weinert LA, Tucker AW, Husna AU, Wileman TM, N FH, Hoa NT, Parkhill J, Maskell DJ, Blackall PJ, Jennings MP. 2018. *Streptococcus suis* contains multiple phase-variable methyltransferases that show a discrete lineage distribution. Nucleic Acids Res doi: 10.1093/nar/gky913:10.1093/nar/gky913.

27. Haigh RD, Crawford LA, Ralph JD, Wanford JJ, Vartoukian SR, Hijazi K, Wade W, Oggioni MR. 2017. Draft Whole-Genome Sequences of Periodontal Pathobionts *Porphyromonas gingivalis, Prevotella intermedia*, and *Tannerella forsythia* Contain Phase-Variable Restriction-Modification Systems. Genome Announcements 5:e01229–17.

28. Qi J, Luo H, Hao B. 2004. CVTree: a phylogenetic tree reconstruction tool based on whole genomes. Nucleic Acids Res 32:W45–7.

29. Xu Z, Hao B. 2009. CVTree update: a newly designed phylogenetic study platform using composition vectors and whole genomes. Nucleic Acids Res 37:W174–8.

30. Qi J, Wang B, Hao BI. 2004. Whole proteome prokaryote phylogeny without sequence alignment: a K-string composition approach. J Mol Evol 58:1–11.

31. Clark TA, Murray IA, Morgan RD, Kislyuk AO, Spittle KE, Boitano M, Fomenkov A, Roberts RJ, Korlach J. 2012. Characterization of DNA methyltransferase specificities using single-molecule, real-time DNA sequencing. Nucleic Acids Res 40:e29.

32. Murray IA, Clark TA, Morgan RD, Boitano M, Anton BP, Luong K, Fomenkov A, Turner SW, Korlach J, Roberts RJ. 2012. The methylomes of six bacteria. Nucleic Acids Res 40:11450–11462.

33. Roberts RJ, Vincze T, Posfai J, Macelis D. 2015. REBASE-a database for DNA restriction and modification: enzymes, genes and genomes. Nucleic Acids Res 43:D298–D299.

34. Guérillot R, Kostoulias X, Donovan L, Li L, Carter GP, Hachani A, Vandelannoote K, Giulieri S, Monk IR, Kunimoto M, Starrs L, Burgio G, Seemann T, Peleg AY, Stinear TP, Howden BP. 2019. Unstable chromosome rearrangements in Staphylococcus aureus cause phenotype switching associated with persistent infections. Proceedings of the National Academy of Sciences of the United States of America 116:20135–20140.

35. Umarov RK, Solovyev VV. 2017. Recognition of prokaryotic and eukaryotic promoters using convolutional deep learning neural networks. PLOS ONE 12:e0171410.

36. de Jong A, Pietersma H, Cordes M, Kuipers OP, Kok J. 2012. PePPER: a webserver for prediction of prokaryote promoter elements and regulons. BMC Genomics 13:299.

37. Fagerlund A, Langsrud S, Schirmer BC, Moretro T, Heir E. 2016. Genome Analysis of Listeria monocytogenes Sequence Type 8 Strains Persisting in Salmon and Poultry Processing Environments and Comparison with Related Strains. PLoS One 11:e0151117.

38. Atack JM, Guo C, Yang L, Zhou Y, Jennings MP. 2020. DNA sequence repeats identify numerous Type I restriction-modification systems that are potential epigenetic regulators controlling phase-variable regulons; phasevarions. FASEB J 34:1038–1051.

